# Replication of alphaviruses requires a pseudoknot that involves the polyA tail

**DOI:** 10.1101/2022.06.20.496828

**Authors:** René CL Olsthoorn

**Affiliations:** Leiden Institute of Chemistry, Leiden University, P.O. Box 9502, 2300 RA Leiden, The Netherlands

## Abstract

Alphaviruses, such as Sindbis virus and Chikungunya virus, are RNA viruses with a positive sense single-stranded RNA genome that infect various vertebrates including humans. A conserved sequence element (CSE) of ∼19 nucleotides (nts) in the 3’ non-coding region is important for replication. Despite extensive mutational analysis of the CSE no comprehensive model of this element exists to date. Here it is shown that the CSE can form an RNA pseudoknot with part of the polyA tail and is similar to the human telomerase peudoknot with which it shares 17 nts. Mutants that alter the stability of the pseudoknot were investigated in the context of a replicon of Sindbis virus and by native gel electrophoresis. These studies reveal that the pseudoknot is required for virus replication and is stabilized by UAU base triples. The new model is discussed in relation to previous data on Sindbis virus mutants and revertants lacking (part of) the CSE.

## Introduction

For RNA viruses it is known that the secondary and tertiary structure of their RNA genome plays an important role in many aspects of their life cycle. RNA viruses have evolved a large variety of RNA structures that are required for replication of their genome (Ahlquist, 1992; You & Rice 2008; Liu et al 2009; 2020; Pflug et al. 2014) regulating their translation (Brierley et al., 1989; Miras et al., 2017; Jaafar & Kieft, 2019; Bhatt et al., 2021) packaging progeny RNA (Carey et al., 1983; Turner et al.,1988; Ding et al., 2020; Frolova et al., 1997; Masters, 2019), or derailing host defense mechanisms (Pijlman et al., 2008; Akiyama et al., 2016; Flobinus et al., 2019; Dilweg et al., 2019). In many instances the 3’- end of plus strand RNA viruses was found to fold into a pseudoknot structure that plays an essential role in the synthesis of a minus-strand copy of the genomic RNA (Rietveld et al., 1983; Pilipenko et al., 1992; Kolk et al., 1998; Tsai et al., 1999; Pogany et al., 2003), sometimes in direct equilibrium with an alternate conformation consisting of stem-loop structures that performs a different function in the viral life cycle (Olsthoorn et al., 1999; Dreher 2009).

Alphaviruses are RNA viruses with a positive-sense single-stranded RNA genome that are found nearly worldwide and include Sindbis virus (SINV), Chikungunya virus (CHIKV) and Eastern and Western equine encephalitis viruses. They are transmitted by mosquitos mostly arthritis and encephalitis. SINV infection of humans mostly os and infect various vertebrates such as humans, horses, rodents, fish, birds as well as invertebrates, causing mostly arthritis and encephalitis. SINV infection of humans mostly occurs in Northern Europe where it is endemic (https://www.ecdc.europa.eu/en/sindbis-fever/facts). CHIKV is endemic in tropical Africa and Asia, though outbreaks have occurred in Italy (Charrel et al. 2007), India (Ramachandran et al. 2012) and more recently in Central America and the Caribbean (Morrison 2014). Since then, CHIKV has been isolated on many continents including America, Europe and Australia (Levi & Vignuzzi, 2019). Presently, no antivirals or vaccines against alphaviruses are commercially available.

The genome of alphaviruses is 11-12 kilobases in length and possesses a 5’ cap structure and a 3’polyA tail. The 5’ two-thirds of the genome encode the polymerase proteins. The capsid protein is encoded by a subgenomic mRNA (26S RNA). Among the studied cis-acting signals are elements needed for 26S RNA synthesis (Levis et al., 1990), encapsidation (Kim et al. 2011), and replication (Hyde et al. 2015; Kendall et al., 2019). Two important elements required for production of minus strand RNA have previously been identified: an RNA structure in the 5’ untranslated region (UTR) (Frolov et al. 2001; Niesters & Strauss, 1990) and a conserved sequence element (CSE) in the 3’-UTR of ∼19 nucleotides (nts) [Levis et al. 1986]. The 5’UTR hairpin has been shown to bind to the viral RNA-dependent RNA polymerase (RdRp) [Frolov et al. 2001]. The role of the CSE in minus-strand synthesis has been investigated in vivo (Kuhn et al. 1990; Raju et al. 1999) and in vitro (Hardy & Rice, 2005; Hardy, 2006). Whereas single substitutions in the CSE can be lethal to the virus, several large deletions in the CSE are tolerated and even complete removal of the CSE can result in viable virus (Kuhn et al. 1990; Raju et al. 1999). In vitro assays have indicated that a polyA tail of at least 11-12 residues is required to get some minus strand synthesis, but that 25 As allow maximal synthesis (Hardy & Rice, 2005). Single substitutions in the CSE are in general detrimental for RNA synthesis although insertions are often tolerated. Altogether the mutational analyses have not led to a comprehensive model for the structure of the CSE in alphaviruses.

Here the possibility of pseudoknot formation by the CSE and part of the polyA tail in alphaviruses is investigated. Using Sindbis virus replicons harboring GFP or luciferase reporter genes it is demonstrated that this pseudoknot structure is essential for viral replication. In addition native gel electrophoresis shows that the pseudoknot is stabilized by UAU base triples by analogy to the telomerase RNA pseudoknot. The new model is discussed in relation to previous data on viable SINV mutants and revertants lacking (part of) the CSE.

## Results

### Conserved Sequence Element (CSE) can fold into a pseudoknot

The CSE has been found to be strongly conserved among alphaviruses (Pfeffer et al., 1998), however the significant variation in its length, 16-21 nts, suggests that the CSE may not function as a linear sequence. By including the polyA tail in the alignment it becomes clear that there is a putative interaction between nucleotides at the 5’ end of CSE and its 3’-end plus several As from the polyA tail (highlighted in green in Figure 1A.) This interaction is supported by covariations: the G_-14_-C_-1_ base pair (bp) in SINV is a UA bp in Sagiyama (SAG), Getah (GET), Ross River (RR), Barmah Forest (BF) viruses and in two Eastern equine encephalitis virus (EEEV) isolates. Vice versa the U_-15_-A_+1_ bp in SINV is a GC bp in SAG, GET, RR, BF, Middelburg (MIDV), Bebaru, Una, and in fish alphaviruses. Likewise, the U_-18_- A_+4_ bp is replaced by an AU bp in several fish alphaviruses. The resulting hairpin is illustrated for several alphaviruses in Fig. 1B. The loop of 10-14 nts includes two stretches of U residues that can form a pseudoknot with the polyA tail as shown in Fig. 1C.

**Fig. 1.**
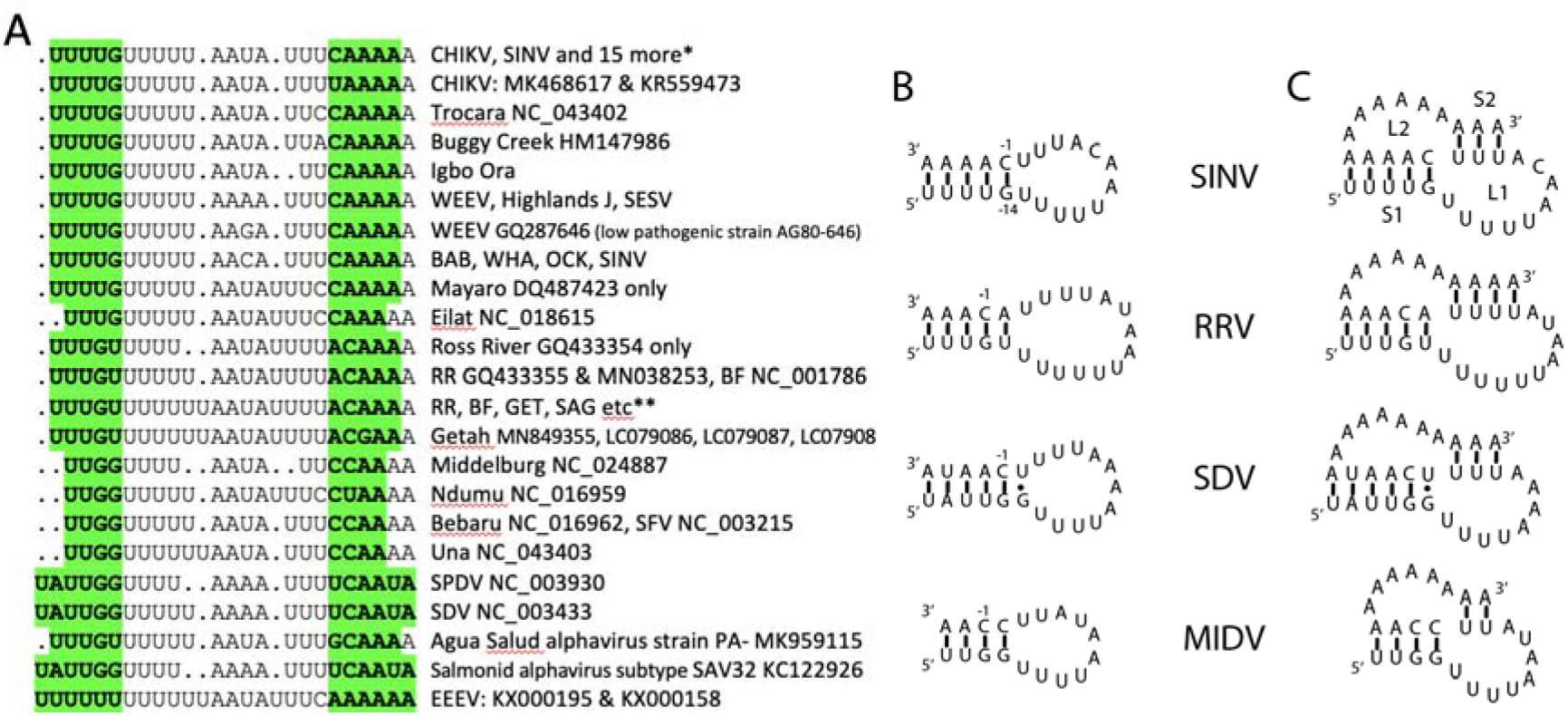
A. Alignment of CSE from all known alphaviruses to date for which the 3’-end is complete. * CHIKV NC_004162, Mayaro NC_003417, VEEV Venezuelan Equine Encephalitis virus NC_001449. Aura NC_003900, EEEV Eastern Equine Encephalitis virus NC_003899, Madariaga NC_023812, Mucambo NC_038672, Mwinilunga LC361437, Tai Forest NC_032681, Fort Morgan NC_013528, Cabassou NC_038670, Rio Negro NC_038674, Mosso das Pedras NC_038857, Pixuna NC_038673, Tonate NC_038675, Everglades NC_038671, O’nyong nyong NC_001512, Igbo Ora AF079457, SINV NC_001547, WEEV Western equine encephalitis virus NC_003908, Highlands J NC_012561, SESV Southern elephant seal virus HM147990, Babanki HM147984, Whataroa NC_016961, Ockelbo M69205, SINV MF589985, MF409177. ** Ross River MH987781 plus others, Barmah Forest MN115377 plus others, Alphavirus M1 EF011023, Getah NC_006558, Sagiyama AB032553. B. Putative hairpin structures formed by the CSE in diverse alphaviruses. SDV: Sleeping disease virus. C. Putative pseudoknot structures involving the polyA tail.

The number of A residues that is required to allow pseudoknot formation in SINV is estimated to be ∼12: 4 As in stem S1 and 3 As in stem S2 plus another 5-6 As in L2, the loop that crosses the minor groove of S1 (Fig. 1C). The minor groove of a 5-bp stem is ∼25 Ångstroms (Pleij et al. 1985) which would require at least 5 As to span the minor groove. Six As are chosen here as one adenosine is predicted to pair with a uracil in L1 (see below). The number of 12 As fits well with the experimentally determined minimum number of 11-12 As that are required for minus-strand synthesis in vitro using SINV RNA (Hardy & Rice, 2005).

### Minimum length of polyA tail required for replication

To determine the minimal length of the polyA tail required for in vivo replication, mutations were introduced via PCR using a SINV replicon expressing GFP (Bredenbeek et al., 1993). In vitro transcribed and capped mRNAs were transfected into BHK21 cells and the number of GFP positive cells was counted ∼20 hr post-transfection. A construct carrying a 29-nt polyA tail (SinA29 or SinA for short) served as a wild type control against which all further constructs were tested. Shortening the length of the polyA tail to 20 As had a minor effect on the number of GFP-positive cells, but at 14, 11 and 8 As an approximately 3-fold reduction in the number of GFP-positive cells was observed (Fig. 2). This reduction was even more dramatic with 4 As (∼1%) and without a polyA tail virtually no GFP-positive cells were visible. This suggests that below 8 As a critical length is reached that severely impairs replication of the virus.

**Fig. 2.**
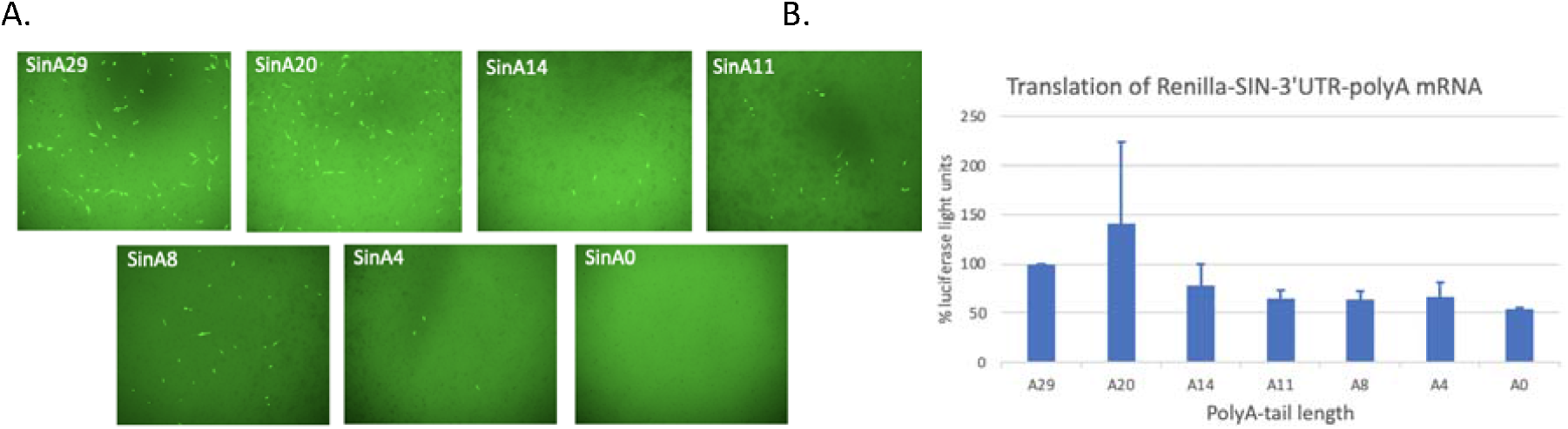
Effect of polyA tail length on replication and translation A. GFP expression in BHK21J cells transfected with SINV replicon RNAs harboring different sizes of the polyA tail. Images show a representative area of a 48-wells plate and are intentionally overexposed to show the density of cells present in this area of the plate. B. Luciferase expression in BHK21J cells transfected with *Renilla* luciferase mRNAs harboring the 3’-UTR of SINV with the indicated length of the polyA tail. Error bars indicate standard deviation of 2 independent experiments.

To investigate to what extent the length of the polyA tail affects the translation of SINV RNA, transfections with a *Renilla* luciferase mRNA harboring the 3’-UTR of SINV with different lengths of the polyA tail were performed. Shortening the polyA tail from 29 to 20 residues appeared to increase translation, while further truncation to 14 and 11 As reduced translation to 78% and 65%, respectively (Fig. 2B). Interestingly, the constructs with 4 or no As still supported a substantial level of translation (54-67%). This suggests that the low or absent replication of SINV RNAs with 4 or less As is predominantly due to a defect in RNA synthesis by the inability to form the pseudoknot, rather than in translation.

### Base pairing in stem 1 contributes to replication

To investigate whether the individual stems of the proposed pseudoknot are required for replication, base pairs were disrupted and restored by secondary mutations. Although in these assays the relative replication levels of mutants with respect to wt were calculated by comparing the number of GFP-positive cells, these levels were treated in a semi-quantitative manner based on the indicated five categories (Fig. 3, inset). As G_-14_ and C_-1_ are strongly conserved nts in the CSE and contribute to the formation of stem S1 they were subjected to mutational analysis first. Disruption of the G_-14_-C_-1_ bp led to a significant decrease in replication (Fig. 3, SinB1), whereas restoring base pairing by the introduction of an AU bp increased replication with respect to the AC mismatch (Fig. 3, SinB2). The AU bp was still worse than wt but this may be due to its lower thermodynamic stability. Disruption of the U_-15_-A_+1_ bp also reduced replication, but this could be restored to wt level by a GC bp (Fig. 3, SinC1 and SinC2). Likewise, disruption of two UA bps in S1, thereby essentially disrupting stem 1, resulted in a low level of replication but restoring stem 1 by formation of AU bps resulted in a significant increase in replication (Fig. 3, SinD1 & SinD2). Finally, replacing S1 with GC-richer stems did reduce replication (Fig. 3, SinE2, SinF, SinG) but also here mismatches reduced replication even more (Fig. 3, SinE1).

**Fig. 3.**
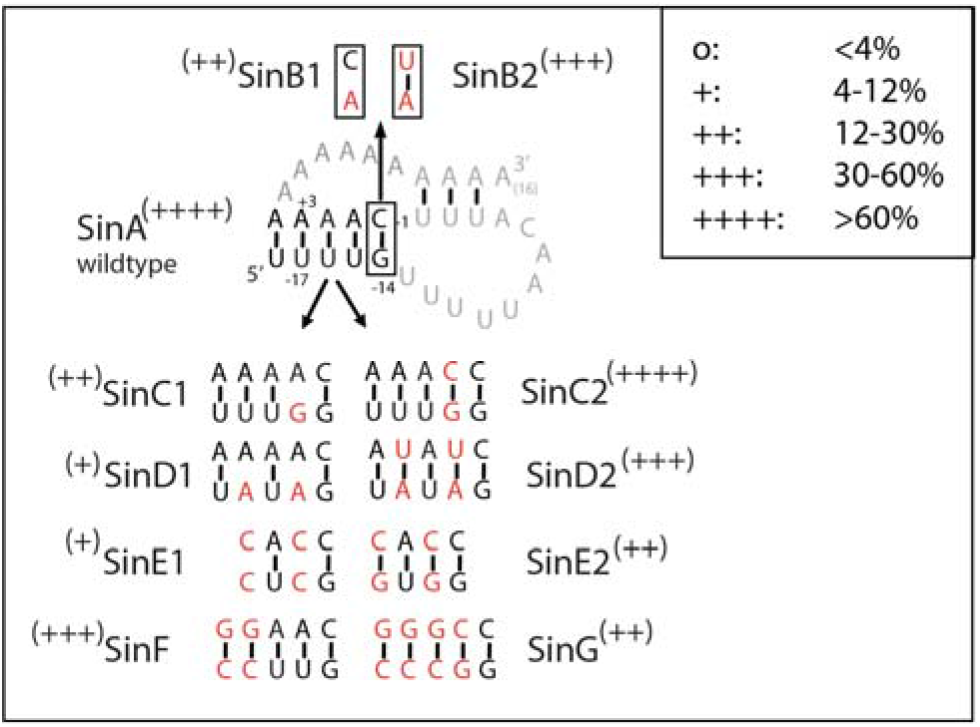
Role of S1 length and stability for replication. Percentages were calculated from at least two independent transfections using wild type SinA as 100% control.

From these results it can be concluded that nts U_-18_ to G_-14_ of the SINV CSE do not function as a linear sequence but are base paired to C_-1_ and four As of the polyA tail thereby forming a stem whose thermodynamic stability can only be varied within certain limits.

### Base pairs in stem 2 are involved in base triples

Stem 2 is predicted to consist of two (e.g. MIDV), three (e.g. CHIKV, SINV, SDV), or four bps (e.g. RRV) in the different alphaviruses (Fig. 1A). Insertion of an additional U residue in SINV PK, thereby creating a 4-bp stem 2, had no significant effect on replication (Fig. 4, SinH). Deletion of one U resulting in a 2-bp stem 2, on the other hand, significantly decreased replicon activity (Fig. 4, SinI). In the context of the MIDV CSE two UA bps in S2 were less detrimental for replication (Fig. 4, inset). This is probably due to the higher stability of S1 as the C to U change in loop L1 has no significant effect on replication (Kuhn et al., 1990; Hardy & Rice, 2005; Fig. S2). Disruption of the U_-2_A_+11_ bp (SinJ) or the U_-3_A_+12_ bp (SinK1), thereby creating a CA mismatch in stem 2, resulted in a reduced or slightly reduced level of replication (Fig. 4). Repairing the CA mismatch in SinK1 by creating a CG match did not increase activity but decreased activity even more (Fig. 4, SinK2). Although this result would suggest that nucleotides at position -3 and +12 are not forming a base pair it is also possible that the UA bps in stem 2 are involved in base triples with U residues in L1 as has been shown for example for the telomerase pseudoknot, Kaposi’s sarcoma-associated herpesvirus (KSHV) polyadenylated nuclear RNA and MALAT-1 non-coding RNA (Theimer et al., 2005; Mitton-Fry et al., 2010; Brown et al. 2012). In those studies, UAU base triples could be replaced with isosteric CGC triples. Such a strategy would also tell which U residues are involved in which base triples in SINV because beforehand it is difficult to predict whether SINV adopts the KSHV or the telomerase-like PK (Fig. 5).

**Fig. 4.**
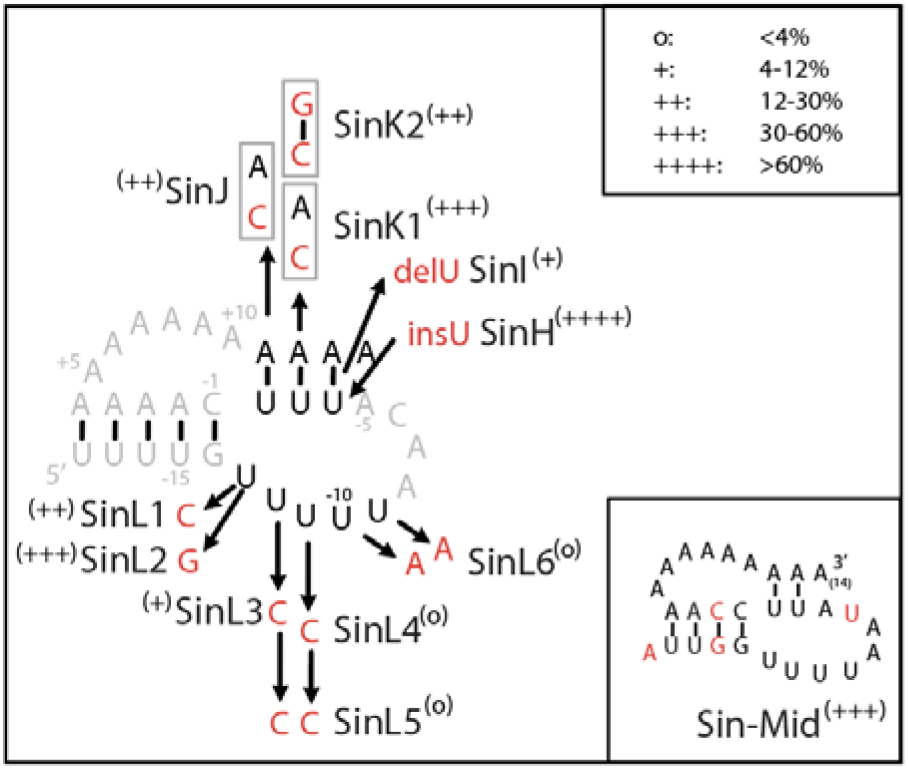
Stem 2 and Loop 1 mutants. Percentages were calculated from at least two independent transfections using wild type SinA as 100% control.

**Fig. 5.**
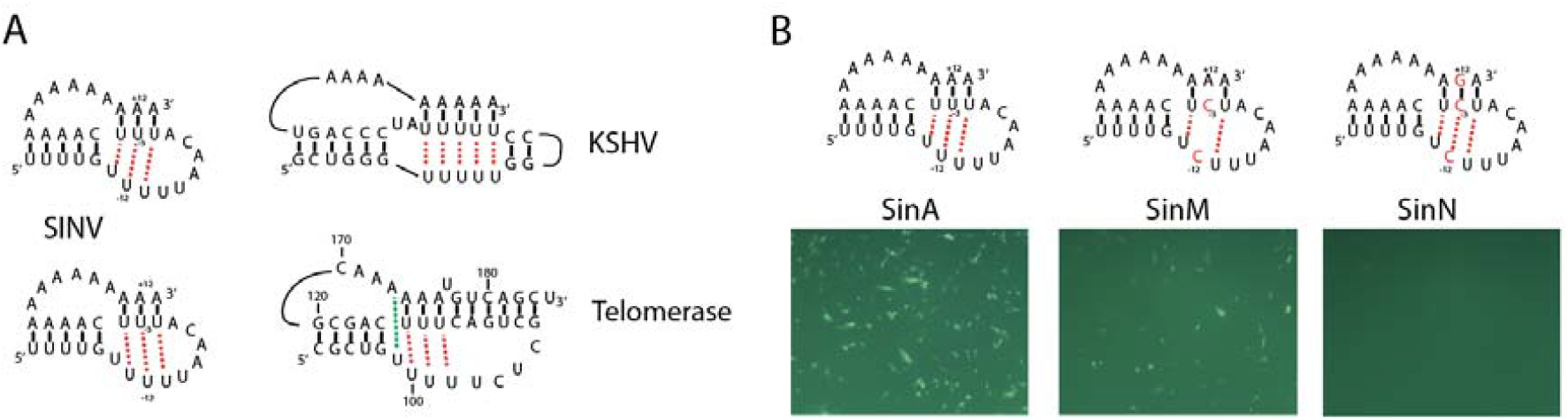
Putative base triples in the SINV pseudoknot, adopting either a KSHV-like conformation (top) or a human telomerase pseudoknot-like conformation (bottom). Red dotted lines indicate Hoogsteen base pairing of U with an AU base pair; green dotted line indicates a Hoogsteen-Watson Crick base pair in the telomerase pseudoknot. B. Testing the KSHV-like pseudoknot conformation by mutating a putative base triple between U_-3_, A_+13_ and U_-12_ using the SINV replicon.

To find out whether the SINV pseudoknot adopted the KSHV-like structure mutations were made in a putative base triple between U_-3_, A_+13_ and U_-12_ (Fig. 5B). As illustrated by the number of GFP-positive cells replication of SinM was a lot worse than wt (SinA). However, replication of SinN which possessed a putative CGC triple was even worse (Fig. 5B). This could mean that the SINV PK does not form base triples at all or that it adopts a telomerase-like PK with base triples in another register. To investigate the latter possibility the putative triple formed by U_-3_A_+12_U_-11_ was disrupted by changing the Us to Cs. This basically abolished replication (Fig. 6A, SinO). However, restoring the base triple with the isosteric CGC triple effectively restored replication (Fig. 6A, SinP). This strongly suggested that the SINV PK favors a telomerase-like PK conformation. Although CGC triples require protonation of one of the cytosines, which in vitro can be achieved by lowering the pH, CGC triples are naturally present in a variety of RNAs (Devi et al., 2015) suggesting that in vivo protonation does occur.

**Fig. 6.**
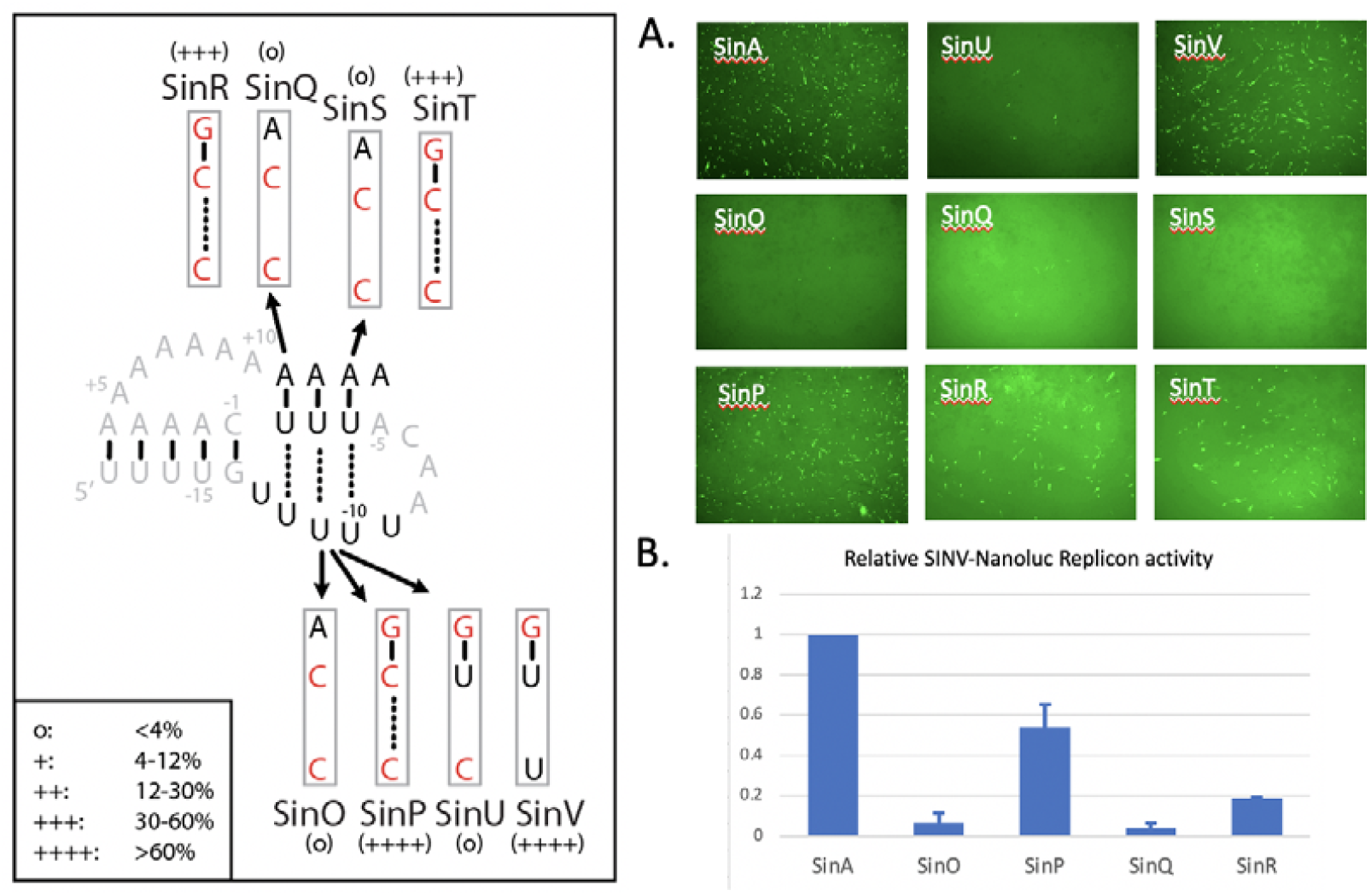
Disruption and restoration of base triples in SINV pseudoknot. A. GFP expression in BHK21J cells transfected with SINV replicons carrying the indicated changes in the pseudoknot. B. Relative nanoluciferase expression of BHK21J cells transfected with SINV-Nanoluc replicons. Error bars indicate standard deviation of 2 experiments.

To investigate whether the other two UA pairs are also stabilized by a base triple they were replaced by CGC triples as well. As can be seen in Figure 6 mutation of U_-2_ and U_-12_ (SinQ) or U_-4_ and U_-10_ (SinS) abolished replication while introduction of CGC triples (SinR and SinT) was able to restore replication significantly, although not to wt levels. These results were largely confirmed by using SINV replicons in which the GPF gene was replaced by a nanoluciferase gene (Fig. 6B).

Additional mutations of the middle UAU triple confirmed that only UAU or CGC combinations are fully functional at this position in the pseudoknot. Mutation of U_-11_ to C was by itself sufficient to virtually knock out replication completely (Fig. 4, SinL4) and a UGC combination was also quite detrimental (Fig. 6, SinU). Interestingly replacing A_+12_ by a G (Fig. 6, SinV) had hardly any effect, even though it might affect the triple interaction. Apparently, this change can be easily accommodated in the structure by a shift in register of the polyA tail, thereby decreasing or enlarging the size of the A-rich loop.

The importance of U_-10_ U_-11_ U_-12_ residues was further emphasized by mutants U_-12_C (Fig. 4, SinL3), U_-12_C+U_-11_C (SinL5) and U_-10_A+U_-9_A (SinL6) which all resulted zero or close to zero replication (Fig. 4). U_-13_ on the other hand is, by analogy with U_99_ in the telomerase pseudoknot (Fig. 5), probably involved in a Hoogsteen base pair with A_+10_. Replacing U_-13_ by a C (Fig. 4, SinL1) reduced replication more than replacing U_-13_ by a G (SinL2) which in principle can form a similar Hoogsteen pair with A_+10_. These results provide additional support for a telomerase pseudoknot-like structure of the SINV 3’- end.

### Native gel-electrophoresis

To get more insight into structural effects caused by mutations in the SINV pseudoknot native gel-electrophoresis with short RNA oligonucleotides was performed. Three SINV-like RNAs SINA, SINB and SINC of 33 nts were studied. These constructs carried two GC base-pairs substitutions at the 5’ proximal end of S1 to stabilize the pseudoknot as initially the wt sequence of the SINV pseudoknot adopted too many conformations to be useful for native PAGE (data not shown). SINA resembles the pseudoknot of the SinF construct which was replicating close to wt level (Fig. 3). As can be seen in figure 7, SINA migrated faster than SINB indicative of a more compact structure. Introduction of a CGC triple (SINC) however did not result in a wt like migration, in fact its migration was even less than that of SINB. As the C in CGC triples needs to be protonated to form two hydrogen bonds with guanine, and the running buffer and gel had a pH of 8.3 the samples were re-run under acidic conditions. Interestingly, at pH 5.5 SINC now migrated faster than SINB and as fast as SINA indicating that protonation indeed promotes formation of the CGC triple.

**Fig. 7.**
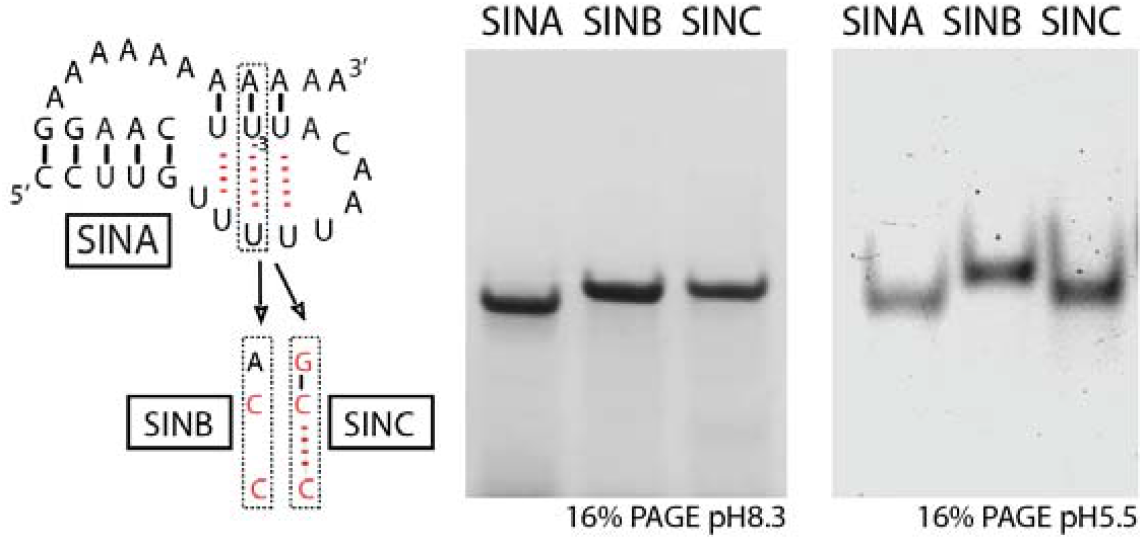
Native gel-electrophoresis at pH 8.3 and 5.5 of SINV pseudoknot RNAs with wild type (SINA) or mutant U_-3_A_+12_U_-11_ base triples (SINB, SINC). RNAs were visualized by Stains-All (left gel) or Ethidium bromide (right gel).

Similar results were obtained with the shorter MIDV pseudoknot (27 nt). Figure 8 shows that at pH 8.3 MidA migrated faster than mutants which had single (MidF) or double (MidB) U to C changes and also faster than MidC with the CGC triple. Again, at pH5.5 MidC catched up with MidA while MidB and F did not, indicating that in MidC the CGC triple is stabilized under acidic conditions. MidD and MidE are shorter variants of MidA that possess a polyA tail of 6 and 9 As resp. As MidE migrated only slightly faster than MidA, it may be assumed that it is still forming the pseudoknot structure. In MidD the number of As is too small to adopt a stable pseudoknot so it probably forms only stem S1 but this remains speculative.

**Fig. 8.**
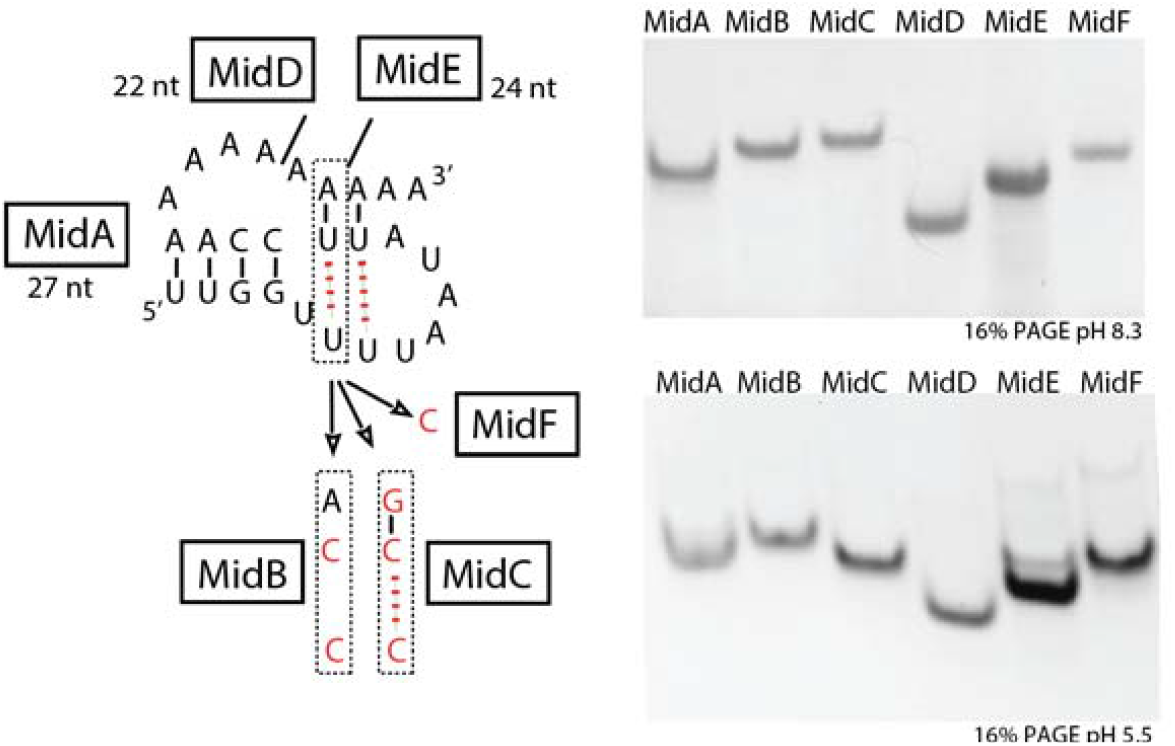
Native gel-electrophoresis at pH 8.3 and 5.5 of MIDV pseudoknot RNAs with wild type (MidA) or mutant U_-3_A_+12_U_-11_ base triples (MidB, C, F), or with shorter polyA tails (MidD and MidE). RNAs were visualized by Stains-All (upper gel) or Ethidium bromide (lower gel).

## Discussion

The data presented in this work strongly suggest that the SINV CSE is not functioning as a linear sequence but as a pseudoknot involving part of the polyA tail. The presence of similar pseudoknot in all alphavirus RNAs is supported by covariations in the base pair composition of stem 1. As stem 2 of the pseudoknot consists solely of base pairs that involve the polyA tail no covariations are found here; the existence of stem 2 in SINV was proven here by replacing the three predicted UAU base triples with CGC triples. In the majority of other alphaviruses also three UAU triples are predicted although some viruses can form four (Fig. 1, e.g. Ross River, Barmah Forest, Getah and Sagiyama viruses). Four viruses, MIDV, Trocara, Buggy Creek, and Igbo Ora virus, can only form two triples (Fig. 1A) but as shown here for MIDV using native gel-electrophoresis this still allows pseudoknot formation.

As the pseudoknot structure depends on the polyA tail, a certain number of As must be present to allow its formation. It was found here that a minimum number of 8 A residues is sufficient to sustain a substantial level of replication compared to SINV RNA having 29 As. Eight A residues are in principle sufficient to form a pseudoknot if two UA bps in stem 1 are present and L2 consists of only 3 bases instead of 6; as discussed below data exist to support that three base pairs still allow minus-strand synthesis. From in vitro studies it was previously determined that at least 11-12 As are required for minus-strand synthesis of SINV RNA but that 10 or fewer As yield only 3% of minus-strand RNA (Hardy & Rice, 2005). This discrepancy can have multiple causes. The in vitro assays may lack a factor that stabilizes the RNA pseudoknot and/or its interaction with the polymerase and in vitro assays may also be more prone to non-native alternative structures that compete with the pseudoknot structure. In vivo nucleotides can be added by the viral polymerase that eventually leads to a better template for minus-strand synthesis. Long stretches of U and A blocks at the 3’-end of SINV mutants were found in vivo (Raju et al. 1999). These are probably added by the nsP4 subunit or a truncated form of nsP4 of the viral polymerase as shown by Tomar et al. (2006). Whether this activity is present in vitro may depend on purification procedures (Thal et al. 2007).

Reinterpretation of the mutations tested by Hardy & Rice (2005) with the pseudoknot model in mind leads to the following observations: mutations upstream of S1 or in the 5’ distal part of S1 do not have a large effect on replication (56-110%, Fig. 9). With the exception of A_-20_A_-19_-to-UU mutant these mutations reduce the length of S1 to 3 bps but do not disrupt it. The G_-14_A mutation shows 68% activity; even though the same mutation (SinZ) here showed only 22% activity, it still suggests that an AC mismatch near the junction with S2 is not very detrimental. Removal of G_-14_ would lead to C_-1_ bulging out, a situation similar to insertion of a C next to C_-1_, both replicating at 62%. The presence of a base at the junction between two coaxially stacked stems of a pseudoknot is not uncommon (van Batenburg et al. 2000). Insertion of other nts than C is more detrimental: insA (13%) and insG (18%) could base pair with U_-13_, thereby disrupting its Hoogsteen bp with A_+10_, as also shown here to affect replication. insU (15%) would create a longer S2 which in the present study was shown to replicate at 77%. Changes in C_-1_ are more detrimental (G:2.2%, U:8.1%) apart from changing the stability of S1 they potentially trigger the formation of alternative hairpins like A_-25_ACAAaauuUUGUU_-12_ or U_-15_GUUuuuAACA_-5_ (stems indicated in capitals) that may compete with the pseudoknot structure.

**Fig. 9.**
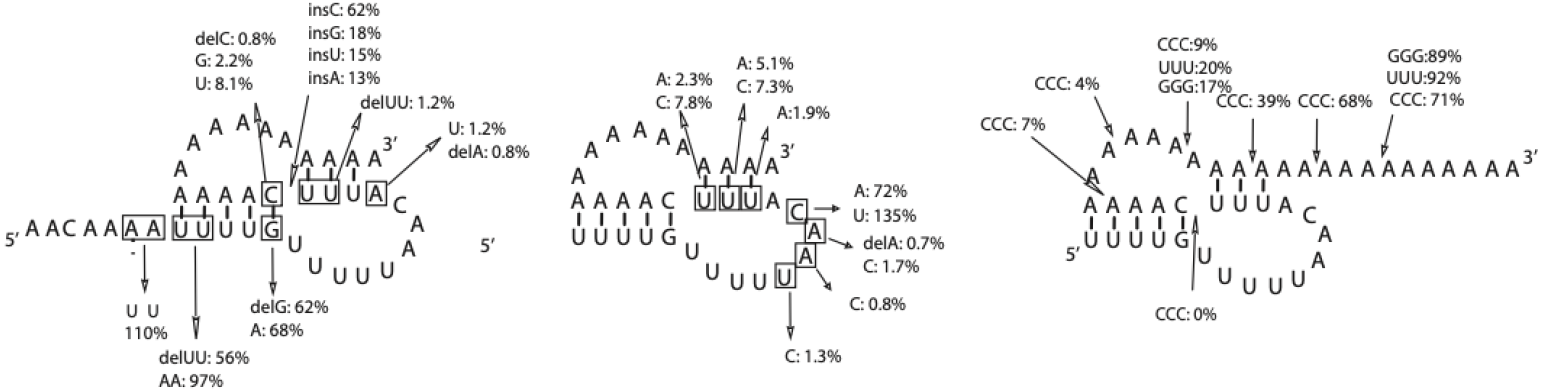
Data from in vitro (-) RNA synthesis of Hardy & Rice (2005) in relation to the pseudoknot structure.

Although deletion of C_-1_ was previously shown to be lethal for SINV replication in vivo (Kuhn et al., 1990) and in vitro (Hardy & Rice, 2005), George & Raju (2000) found that replicating virus could be obtained after deleting C_-1_. The sequence of these replicating mutants showed insertions of six U residues directly upstream of the polyA tail. Furthermore, Hardy & Rice (2005) showed that template activity of the C_-1_ deletion mutant could be gradually restored from 10-19% by inserting one to five U-residues. It should be noted that these additional Us can base pair with As from the polyA tail thereby extending the length of stem S2; this may benefit the stability of the entire pseudoknot compensating for the absence of the G_-14_C_-1_ pair.

Deletion of U_-1_ and U_-2_ (1.2%, Fig. 9 left) disrupts S2 and is therefore hardly viable. Introducing AA mismatches in S2 is more detrimental than AC mismatches (Fig. 9, middle), likely because the latter ones are less destabilizing than AA mismatches. In this study AC mismatches in S2 retained 30-40% activity (Fig. 4, SinJ and SinK1). Changes in the unpaired U_-10_AACA_-6_ sequence were mostly (close to) lethal (Fig. 9, middle). This is a strongly conserved motif in all alphaviruses, except for the C which is not conserved and can be replaced by other nucleotides without major effects. These results were also confirmed in this study although the “lethal” A_-8_C mutant still retained ∼9% activity (Supplementary Fig. S1).

Hardy & Rice (2005) also measured the effect of inserting a row of three C’s at different locations in the polyA tail (Fig. 9, right). Insertions near C_-1_ abolished minus-strand synthesis, likely due to distorting the stability of the pseudoknot and the orientation of stems with respect to each other. Insertion of CCC between A_+3_ and A_+4_ (7%), though still preserving 4 out of 5 base pairs in S1, changes the composition of L2 substantially, putatively disrupting A-minor interactions with S1. A-minor interactions are well-known stabilizers of pseudoknots, including the telomerase pseudoknot (Theimer et al. 2005), and also of RNA triplexes involving the polyA tail (Torabi et al. 2021). Likewise, insertion between A_+6_ and A_+7_ (4%) and A_+9_ and A_+10_ (9%) could affect the A-minor interactions with S1 or disrupt base pairs in S2. Insertions downstream of A_+15_ or A_+18_ had minor effects on minus-strand synthesis, most likely because they do not interfere with the pseudoknot structure.

All together the proposed pseudoknot structure seems to be in agreement with the mutational analysis of the CSE by Hardy & Rice (2005).

Can the pseudoknot model for the 3-CSE explain “Raju’s revertants” that were obtained after transfection of severely debilitated mutants of the CSE? (Raju et al., 1999; George & Raju, 2000; James et al., 2007). Many of these mutants acquired alternating blocks of Us and As that were added to the 3’-end of the mutated CSEs by a cellular or the viral polymerase. Careful inspection of these sequences showed that most of them can form a pseudoknot immediately preceding the polyA-tail that resembles the wt SINV pseudoknot in many aspects including the conserved UAANA motif and ≥2 UAU triples (Supplementary Fig. S2). A few of these alternative pseudoknots were tested in the SINV replicon. Some of these were quite successful in replication, i.e. AA34 and AA35, corresponding to revertants 15.3 and S3-4 replicated at 30-40% of wt level (Supplementary Fig. S3). Replication of five other pseudoknots however was poor with levels not exceeding 3% (Supplementary Fig. S3). The fact that not all revertant sequences tested here replicated efficiently may be due to missing sequence elements that contributed to the survival of Raju’s revertants which carried up to 100 additional nts. In the present assays, only the sequence that could be folded into a pseudoknot was incorporated into the replicon. Nevertheless, the viability of several of these mutants and revertants which adopt very weak pseudoknot structures (Fig. S2) suggests that the pseudoknot or the CSE is not essential for replication. However, the pseudoknot does contribute to viral fitness as wt viruses rapidly outcompeted these mutants and revertants during a co-infection (James et al., 2007).

A similar pseudoknot involving the polyA tail is present at the 3’-end of Bamboo Mosaic Virus (BaMV) RNA (Tsai et al., 1999). BaMV is a plant virus of the Bromoviridae family which is part of the Alphavirus superfamily and is closely related to alphaviruses based on conserved motifs in their RdRps. For BaMV the requirement of the pseudoknot by itself for replication has not been tested since this would require mutation of the polyA sequence as well. Recent data suggest that the BaMV pseudoknot is also stabilized by UAU triples and that is a similar pseudoknot is present in the majority of potexviruses and viruses of the Betaflexiviridae (R.C.L.O, C. Owens, I. Livieratos, manuscript in preparation).

Interestingly, BaMV minus-strand synthesis has been found to start from several sites within the polyA tail with the majority starting at position A_+7_ in L2 (Cheng et al. 2002). This differs from findings with SINV by Hardy (2006) who found that minus-strand synthesis occurs opposite of C_-1_ using in vitro assays. On the other hand (-) RNA isolated in vivo was found to contain at least 60 U residues at the 5’end suggesting that replication initiates within the polyA tail (Sawicki and Gomatos, 1976; Frey and Strauss, 1978). Also, many of the revertants found by Raju and coworkers (Raju et al. 1999; George & Raju, 2000) actually stably maintain A and U insertions downstream of C_-1_ in their genome for many passages. The observation that all-A/U pseudoknots in this study (AA34 and AA35) replicated efficiently also suggests that minus-strand synthesis in vivo starts at a different position than in vitro, possibly owing to different reaction conditions.

The SINV pseudoknot is stabilized by UAU base triples and in this respect shows resemblance to other RNAs like KSHV and telomerase peudoknots. By mutational analysis it was demonstrated that the SINV triples are not in the KSHV-like register but adopt the telomerase-like structure leaving U_-13_ to form a Hoogsteen Watson-Crick base pair with an A in the polyA tail by analogy with the human telomerase PK (Fig. 10). Mutation of this potential Hoogsteen bp in SINV led to a substantial decrease in replication, similar to the effects on the human telomerase PK (Theimer et al. 2005). Replacing U_-13_ by a G was less detrimental than the U_-13_ to C change, probably because G can also form a Hoogsteen pair with A_+10_. The structural similarity with the telomerase PK also suggests that A-minor interactions of the A_+8_ and A_+9_ have a stabilizing role in SINV PK (Fig. 10). Their contribution could not be tested in this work due to the fact that the loop size of L2 is quite flexible so that changing these two particular As can easily be compensated by neighboring As in the polyA tail. Future experiments using for example the SinP mutant as a scaffold in which the loop size of L2 is fixed by the CGC triple would allow for such studies.

**Fig. 10.**
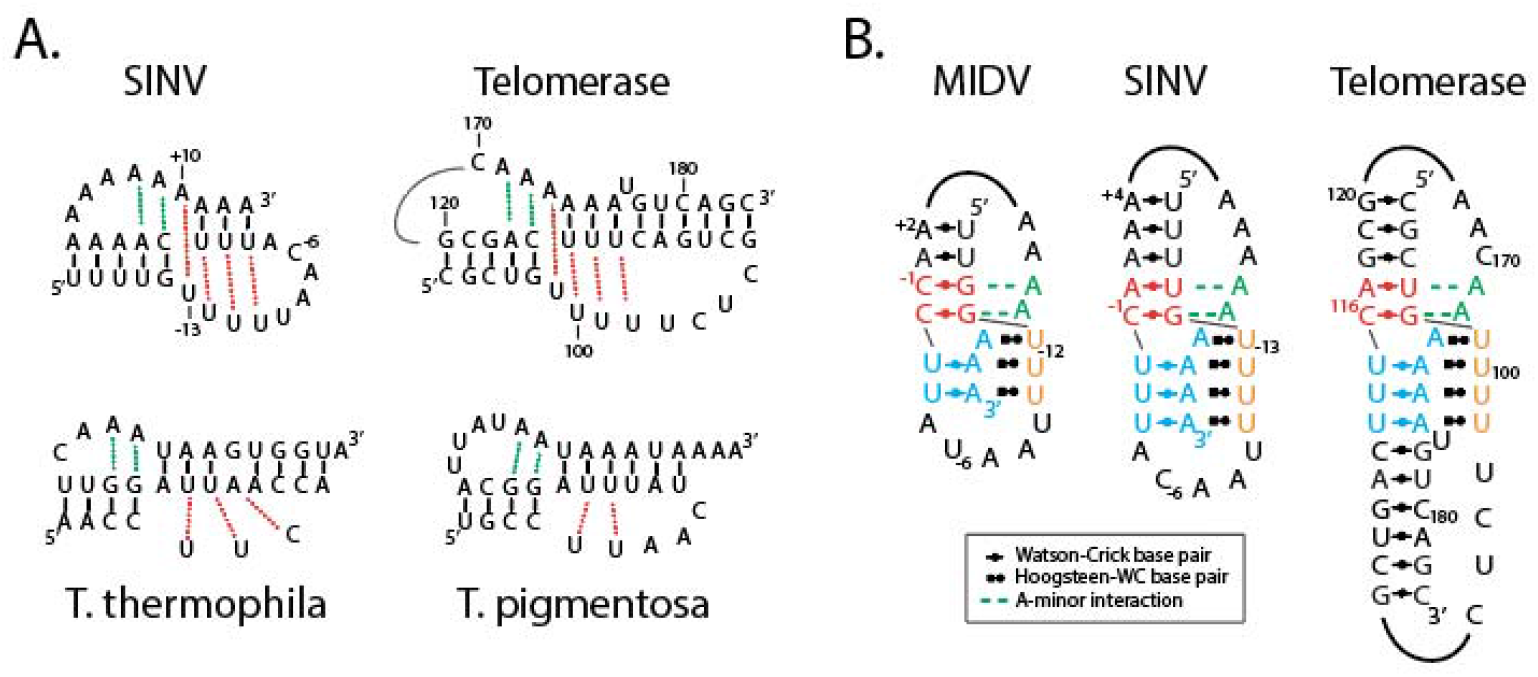
Comparison of alphavirus and telomerase pseudoknots. A. Similarities between SINV and human and tetrahymena telomerase pseudoknots. Dotted green lines indicate A-minor interactions, red dotted lines indicate Hoogsteen Watson-Crick base pairs, black dashes Watson-Crick base pairs. The interactions in *T. pigmentosa* are drawn by analogy to the *T. thermophila* pseudoknot (Cash & Feigon, 2017). B. Detailed models of the MIDV and SINV pseudoknots drawn to resemble the human telomerase pseudoknot (adapted from Theimer et al. 2005).

In terms of stability the SINV PK would be expected to be much weaker than the human telomerase PK even though both structures are active at 37°C. For native gel electrophoresis it was necessary to replace some of the AU pairs in S1 with GC pairs in SINA RNA to obtain well-resolved bands on gel. However, the smaller MIDV pseudoknot was stable under these conditions suggesting that a stem S2 of just 2 AU bps is feasible when stabilized by base triples. Interestingly, some of the telomerase PKs that are present in Tetrahymena species are AU-rich as well (Fig. 10). The *T. thermophila* PK reportedly is not very stable being in an equilibrium with an alternating hairpin structure (Cash & Feigon, 2017). It is likely that the alphavirus PKs are also in equilibrium with a hairpin which may have a different function in the lifecycle of the virus, for instance favoring translation. Pseudoknots at the 3’-end of viral RNAs may fulfil such a role with or without the help of viral or host proteins (Olsthoorn et al. 1999; Dreher 2009). These alternating structures in viral RNAs could be promising targets for small molecule drugs that interfere with viral replication by shifting the equilibrium towards one or the other side. Recently, using high throughput screening approaches small molecule ligands have been identified that either bind the triple helix of MALAT1 non-coding RNA or its hairpin conformation (Donlic et al., 2018; Abulwerdi et al., 2019). Similar HTS approaches directed at the alphavirus pseudoknot could eventually lead to an effective antiviral against encephalitis and Chikungunya.

## Experimental Procedures

### Constructs

Templates for full-length SINV RNA were obtained by PCR on the Sinrep5 plasmid (kindly provided by Dr. P. Bredenbeek, Leiden University Medical Center, The Netherlands) using forward primer SP6SIN and several reverse primers (Supplementary Table 1). PCR conditions were: 95°C for 3 min, then 30 cycles of 94°C 1 min, 55°C 45 sec, 68°C 5 min, followed by 5 min at 72°C. PCR reactions were carried out in a 50 µl volume, containing 400 nM of each oligo, 200 µM dNTPs and 2 units DreamTaq polymerase (Thermofisher) on a BioRad cycler. PCR products were purified by NaAc/EtOH precipitation, washed with 70% EtOH, and after drying dissolved in Milli-Q water.

Sinrep-Nluc was constructed by digestion of Sinrep5 plasmid with Xba and StuI and insertion of an NheI-XbaI (blunted by T4 DNA polymerase) fragment containing the Nluc gene of pNL1.1 (Promega, Benelux). The RLuc-SIN-3UTR plasmid was obtained by insertion of a PCR fragment covering the SINV 3’-UTR into a *Renilla* luciferase reporter plasmid previously described (Girard et al., 2011). The XbaI-XhoI fragment of this vector was replaced with the XbaI-XhoI digested SIN-UTR PCR fragment. Templates for transcription were obtained by PCR using forward primer SP6FLU and reverse primers (Supplementary Table S1).

### Transcription

RiboMAX Large Scale RNA Production Systems (Promega, Benelux) and HiScribe SP6 RNA synthesis kits (New England Biolabs) were used for transcription. Reactions were carried out in 5 or 10 ul volumes containing 100-200 ng of PCR template DNA, ATP, CTP, UTP (10mM each), 2 mM 3’-O-Me-m ^7^G(5’)ppp(5’)G cap structure analog (NEB S1411S), 0.5 mM GTP, 0.5-1 unit of RNAse-inhibitor (RNAsin, Promega), and 0.5-1 ul of enzyme mix. Incubation was done at 37°C for 2.5 hrs. Quality and quantity of transcripts were checked on agarose gels.

### Transfection

100 ng of transcript (as judged by gel electrophoresis) was mixed with Dulbecco’s Modified Eagle’s Medium (DMEM) and 0.5 ul of MessengerMax lipofectamine (Invitrogen) premixed with DMEM and after 20 min incubation at RT added to a well of a 48-wells plate containing BHK21J cells at ∼70% confluency. Cells were grown at 37°C in 5% CO_2_ on growth medium consisting of DMEM supplemented with 10% GlutaMAX (Thermofisher), ampicillin, streptomycin and foetal calf serum, and analyzed 20-24 hrs after transfection. Fluorescent microscopy was performed on an EVOS FL Auto Imaging System (Fisher Scientific). Luciferase activity was measured after lysis of cells using Passive Lysis Buffer (Promega) and transferring the contents to a 96-well plate to which the appropriate substrate was added and assayed using a GloMax multi system (Promega).

### Native PAGE

RNA oligonucleotides were purchased from Merck (Sigma-Aldrich) at 0.05 nmole scale/desalted. 100-200 pmoles of each RNA were loaded onto polyacrylamide gels containing 12% or 16% acrylamide:bisacrylamide (19:1), Tris (40 mM), acetate (20 mM) EDTA (1 mM) pH 8.3, to which 1.5 mM MgAc_2_ was added. Gels were run in TAEM buffer at ∼85 V and 12 mA for ∼4 hrs in a cold room. For acidic PAGE the pH was adjusted by addition of acetic acid to all buffers involved. RNA was visualized by EtBr or Stains-All staining.

## Acknowledgments

I thank all colleagues from the former Plant virology and Genexpress groups of Leiden University and especially Kees Pleij, Sacha Gultyaev and Maarten de Smit for their interest and useful suggestions.

